# Cisplatin Exposure Alters the Effects of Honokiol on Genes Involved in Outer Hair Cell Physiological Function

**DOI:** 10.64898/2026.10.22.761060

**Authors:** Chail Koo, Xiaodong Tan

## Abstract

Outer hair cell (OHC) loss occurs in cisplatin-induced hearing loss (CIHL). We previously reported that honokiol protects against CIHL and mitigates cisplatin-induced transcriptomic changes in OHCs. Here, we re-analyzed OHC RNA-seq to determine whether honokiol differentially affects genes involved in OHC physiological function depending on cisplatin exposure. Twenty-two genes involved in mechanotransduction and hair bundle function, membrane potential and ion homeostasis, electromotility, and efferent regulation were selected. Among 20 genes with available interaction statistics, 16 showed significant Honokiol × Cisplatin interactions, of which 15 were negative. Notably, in the presence of cisplatin, honokiol downregulated several genes associated with mechanotransduction (MET) machinery, including *Lhfpl5, Tmc1, Cib2*, and *Tomt*. Because MET channels have been implicated in cisplatin’s entry into hair cells, these changes raise the possibility that altered MET machinery may contribute to honokiol-mediated protection against CIHL. However, whether these changes decrease cisplatin uptake remains to be determined.

## 1. Introduction

Outer hair cells (OHCs) amplify acoustic signals through a cascade of molecular processes, each involving different proteins (**Fig. 1**). Mechanoelectrical transduction (MET) converts mechanical stimulation in the hair bundles into electrical responses. The MET machinery in mature OHCs involves cadherin 23 (*Cdh23*), protocadherin 15 (*Pcdh15*), lipoma HMGIC fusion partner-like 5 (*Lhfpl5*), transmembrane inner ear (*Tmie*), transmembrane channel-like 1 (*Tmc1*), and calcium and integrin binding family members 2 and 3 (*Cib2* and *Cib3*) (Holt et al., 2021). Hair bundle function involves harmonin (*Ush1c*), USH1G protein (*Ush1g*), myosin VIIA (*Myo7a*), and espin (*Espn*). Transmembrane O-methyltransferase (*Tomt*) is required for MET via TMC1 (Holt et al., 2021). Plasma membrane potential and ion homeostasis are maintained and regulated by voltage-gated potassium channels KCNQ4 (*Kcnq4*) and KCNA10 (*Kcna10*), Na^+^/K^+^-ATPase subunits (*Atp1a1* and *Atp1b1*), and plasma membrane Ca^2+^ ATPase 2 (*Atp2b2*) (Fettiplace & Nam, 2019). Electromotility is mediated by prestin encoded by solute carrier family 26 member 5 (*Slc26a5*) (Zheng et al., 2000). These processes constitute fundamental physiological functions of OHCs and together enable cochlear amplification. OHC function is also regulated by the cholinergic nerve endings of medial olivocochlear (MOC) efferent fibers. Acetylcholine (ACh) activates nicotinic ACh receptor subunits alpha 9 and alpha 10 (*Chrna9* and *Chrna10*) at the basolateral membrane of OHCs (Murthy et al., 2009). Calcium influx activates small- and big-conductance calcium-activated potassium channels, calcium-activated potassium channel subfamily N member 2 (SK2; *Kcnn2*) (Murthy et al., 2009) and subfamily M alpha 1 (BK; *Kcnma1*) (Rohmann, Wersinger, Braude, Pyott, & Fuchs, 2015).

**Figure 1.**
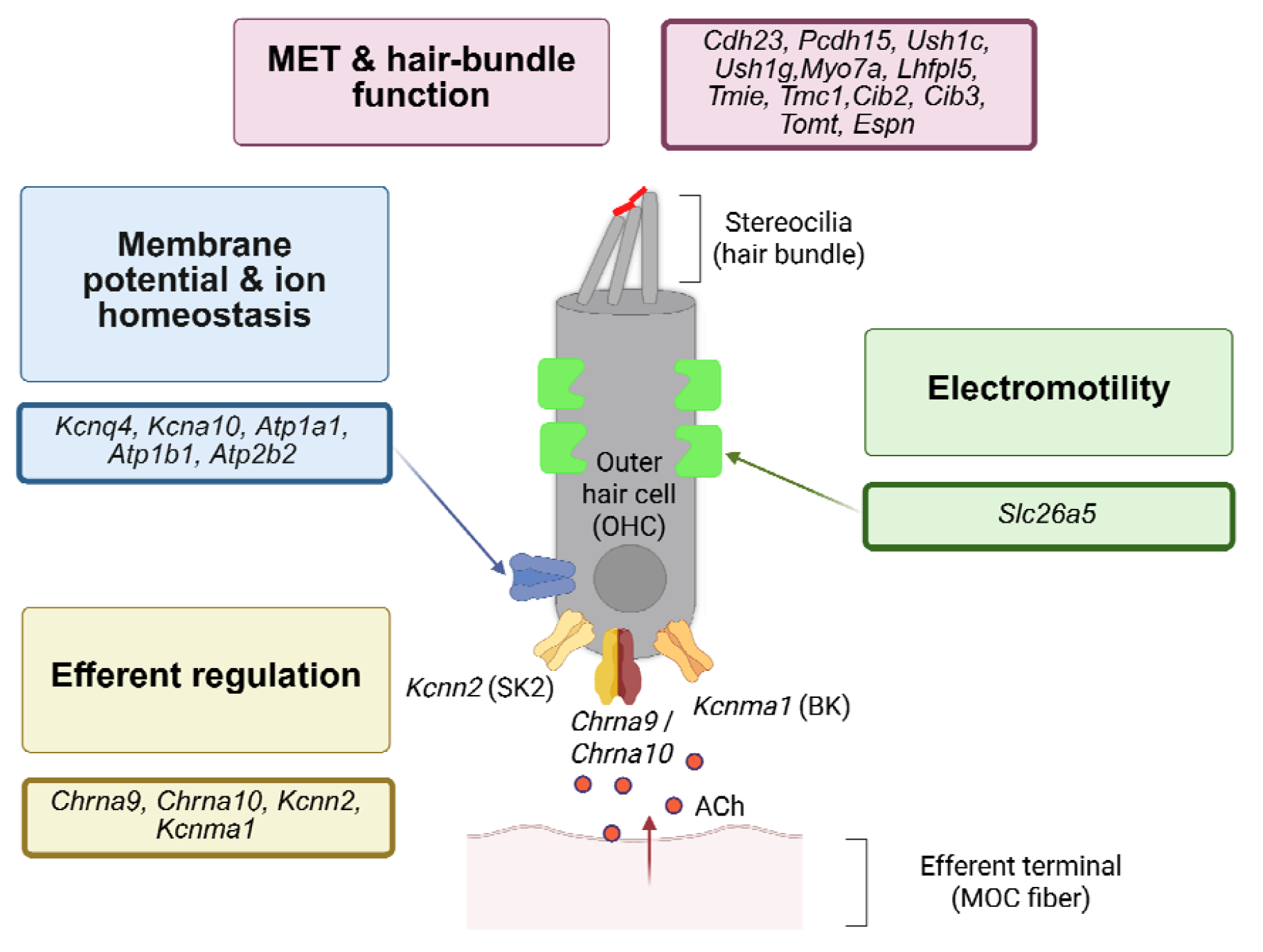
Genes involved in mature outer hair cell physiological function. Mechanoelectrical transduction (MET) is mediated by a multiprotein machinery comprising the MET channel complex (*Tmc1, Lhfpl5, Tmie, Cib2, Cib3*), the extracellular tip link (*Cdh23, Pcdh15*), and associated structural and regulatory proteins (*Tomt, Ush1c, Ush1g, Myo7a, and Espn*). Membrane potential and ion homeostasis are regulated by K^+^ channels (*Kcnq4* and *Kcna10*), Na^+^/K^+^-ATPase (*Atp1a1* and *Atp1b1*), and PMCA2 at the hair bundles (*Atp2b2*). Electromotility is conferred by prestin (*Slc26a5*) in the lateral membrane. Cholinergic efferent signaling from MOC fibers modulates OHC activity through α9α10 nAChRs (*Chrna9* and *Chrna10)* and Ca^2+^-activated K^+^ channels SK2 (*Kcnn2*) and BK (*Kcnma1)*.

Severe OHC loss is observed in cisplatin-induced hearing loss (CIHL), an important clinical problem linked to chemotherapy, affecting half a million cancer survivors globally each year (Dillard et al., 2022). We have reported that honokiol, a multifunctional naturally occurring biphenolic lignan, protects against CIHL in mice. In a recent study with cell-specific bulk transcriptomics of mouse OHCs, we found that honokiol ameliorated cisplatin-induced downregulation of mitochondrial redox genes, and that honokiol exerted transcriptomic changes in different antioxidant pathways depending on whether OHCs were cisplatin-treated or untreated (Koo et al., 2026). However, these pathways only represent cellular responses to stress. Whether cisplatin and honokiol also affect the expression of genes directly related to OHC physiological functions is unclear.

In this brief report, we hypothesized that honokiol-induced transcriptional changes in the genes involved in OHC physiological function would vary depending on whether OHCs had been under cisplatin exposure or not. We re-analyzed the publicly available RNA-seq data (GSE328003) and selected 22 genes based on their established physiological roles; mechanotransduction and hair-bundle function, membrane potential and ion homeostasis, prestin-mediated electromotility, and efferent regulation (**Fig. 1**). Changes in the expression of these genes after cisplatin and honokiol treatment and the interaction between honokiol and cisplatin were studied.

## 2. Materials and methods

RNA-seq data were extracted from the NCBI Gene Expression Omnibus (GSE328003). OHCs were manually aspirated and collected from the cochleae of mice that had received Vehicle, Honokiol, Cisplatin, or Honokiol + Cisplatin (combination) at P21–24. For the combination group, honokiol was given one hour prior to cisplatin (Koo et al., 2026). No new experiments involving animals were carried out in the present study.

For the present study, transcripts from 32 biological samples were re-quantified using Salmon and the *Mus musculus* GRCm39 annotation (Ensembl release 113). Differential expression analysis was carried out using DESeq2. Genes with at least 10 counts in at least three biological samples were kept, resulting in 19,796 genes available for analysis. The DESeq2 model included batch (different library prep time points), honokiol treatment, cisplatin treatment, and the Honokiol × Cisplatin interaction (∼ batch + honokiol + cisplatin + honokiol:cisplatin).

Twenty-two genes involved in OHC function were chosen a priori based on their established roles; mechanotransduction (*Cdh23, Pcdh15, Ush1c, Ush1g, Myo7a, Lhfpl5, Tmie, Tmc1, Cib2, Cib3, Tomt, and Espn*) (Holt et al., 2021), membrane potential and ion homeostasis (*Kcnq4, Kcna10, Atp1a1, Atp1b1, and Atp2b2)* (Kubisch et al., 1999), prestin-mediated electromotility (*Slc26a5*) (Zheng et al., 2000), and efferent regulation (*Chrna9, Chrna10, Kcnn2*, and *Kcnma1*) (Murthy et al., 2009; Rohmann et al., 2015). The selection was intended to reflect the molecular machinery of OHCs, rather than to investigate every gene expressed by OHCs.

The effect of honokiol was studied in the absence or presence of cisplatin. A factorial DESeq2 model was used to test the Honokiol × Cisplatin interaction. The interaction was defined as (combination – cisplatin) – (honokiol – control). This was used to study whether the effect of honokiol was different by cisplatin exposure. P values were adjusted using the Benjamini-Hochberg false discovery rate (FDR) of DESeq2. FDR < 0.05 was deemed statistically significant.

## 3. Results

Of the 22 selected OHC functional genes, complete differential expression statistics were unavailable for *Cib3* and *Pcdh15* (**Supplementary Table 1**); *Cib3* showed low-level expression and did not pass the threshold for differential expression analysis, and the *P*-value of *Pcdh15* was unavailable due to Cook’s distance filtering.

For the remaining 20 OHC functional genes, cisplatin, honokiol, and combination treatment resulted in different expression patterns (**Fig. 2**). For the cisplatin treatment group, most genes did not show statistically significant differences, except for *Kcnn2*, which was significantly upregulated. In contrast, honokiol treatment significantly increased *Lhfpl5, Cib2, Kcna10, Slc26a5*, and *Kcnn2*, while *Myo7a* was significantly decreased. However, in the presence of cisplatin, honokiol significantly decreased *Ush1c, Lhfpl5, Tmc1, Cib2, Tomt, Kcnq4, Kcna10, Atp1a1, Slc26a5, Chrna9, Chrna10, Kcnn2*, and *Kcnma1*.

**Figure 2.**
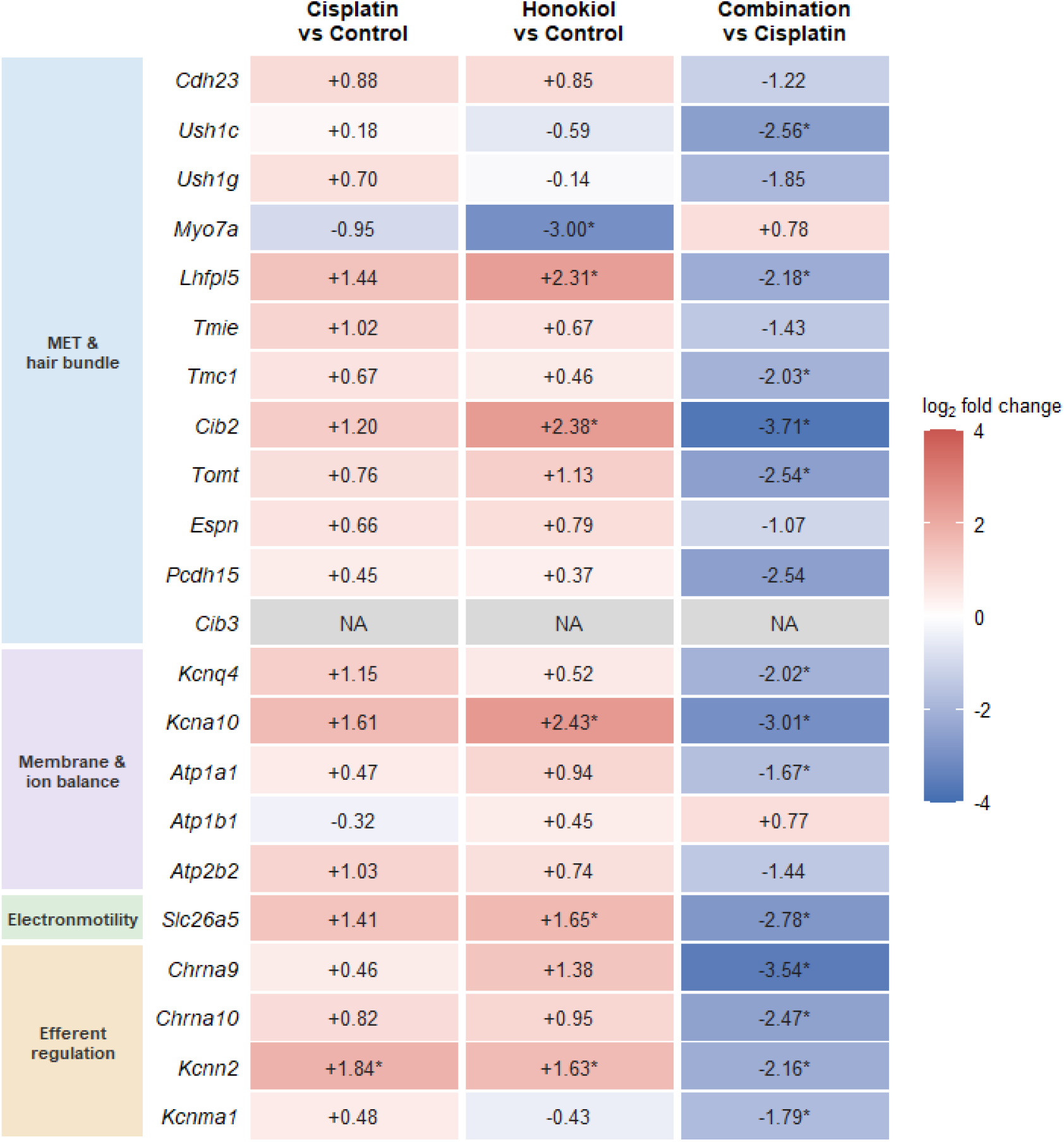
Changes in the expression of genes involved in outer hair cell physiological function after cisplatin and honokiol treatment. Heatmap shows the changes in the expression of selected genes involved in outer hair cell (OHC) physiological function. Genes are grouped by four functions: mechanoelectrical transduction (MET) and hair bundle function, membrane potential and ion homeostasis, electromotility, and efferent regulation. Three treatment comparisons are shown: cisplatin versus control, honokiol versus control, and combination (honokiol + cisplatin) versus cisplatin. In each comparison, red indicates higher gene expression in the first group relative to the second group, while blue indicates lower expression. Values indicate log_2_ fold changes while asterisks (*) indicate statistically significant differences (FDR-adjusted P < 0.05). Grey cells indicate unavailable data. In the combination group, honokiol was administered 1 hour before cisplatin.

In summary, honokiol treatment produced different transcriptional changes in genes related to the physiological function of OHCs depending on whether the mice were untreated or cisplatin treated.

Next, we carried out a formal Honokiol × Cisplatin interaction analysis. Among the 20 genes with available interaction statistics, 16 showed significant interactions. Within the mechanotransduction and hair-bundle module, significant interactions were observed for *Cib2* (log_2_FC = −6.91, FDR = 8.65 × 10− □), *Lhfpl5* (−5.60, FDR = 4.17 × 10^−^ □), *Myo7a* (+3.92, FDR = 2.34 × 10^−^ □), *Tomt* (−4.95, FDR = 0.00237), *Tmc1* (−2.95, FDR = 0.00928), *Cdh23* (−2.61, FDR = 0.0316), *Tmie* (−2.75, FDR = 0.0443), and *Espn* (−2.30, FDR = 0.0473). Significant interactions were also observed for *Kcnq4* (−3.05, FDR = 0.00880), *Kcna10* (−6.87, FDR = 7.82 × 10^−^ □), *Atp1a1* (−3.03, FDR = 0.00169), *Atp2b2* (−2.67, FDR = 0.0199), *Slc26a5* (−5.06, FDR = 8.55 × 10^−^ □), *Chrna9* (−6.25, FDR = 9.51 × 10^−^ □), *Chrna10* (−4.17, FDR = 0.00118), and *Kcnn2* (−4.56, FDR = 2.13 × 10^−^ □). *Ush1c, Ush1g, Atp1b1*, and *Kcnma1* did not show significant interactions. For *Pcdh15*, the interaction log_2_FC was −3.96, but statistical significance was unavailable due to Cook’s distance filtering. *Cib3* was excluded from differential expression analysis because of low expression. Overall, these results show the pattern that the transcriptional effect of honokiol on genes related to OHC physiological functions differs depending on the presence or absence of cisplatin.

## 4. Discussion

In this work, we studied how cisplatin and honokiol affected the expression of genes involved in the physiological functions of OHCs. The unexpected downregulation of OHC functional genes after combination treatment has an important implication for protection against cisplatin. Previous research have shown that MET channels contribute to cisplatin uptake and increase susceptibility to cisplatin ototoxicity (Kitcher et al., 2019; Li, Liu, Kaefer, Youssef, & Zhao, 2022). Genetic disruption of mechanotransduction partially protected hair cells against cisplatin (Li et al., 2022), while pharmacologically blocking MET channel gave protection against cisplatin and aminoglycoside ototoxicity (Kenyon et al., 2021; Kitcher et al., 2019). In the present study, honokiol in the cisplatin presence significantly decreased the genes related to MET machinery in OHCs; *Lhfpl5, Tmc1, Cib2*, and *Tomt*. Interestingly, our previous study showed that honokiol pre-treatment increased OHC survival from 75.3% in cisplatin-treated mice to 88.8% in the combination group (Koo et al., 2026). These findings suggest that the decreased expression of the MET machinery related genes, along with other genes involved in OHC physiological functions, may represent the underlying protective mechanisms by which OHCs respond to harmful insults such as ototoxic drugs.

However, the present study does not establish a causal relationship between gene downregulation and functional suppression. Further studies examining the protein expression and MET current changes in OHCs will help understand this causality.

Interestingly, cisplatin alone had little effect on the expression of the OHC functional genes in this study. Among the 20 genes with available expression profiles, only *Kcnn2* was significantly changed by cisplatin compared to controls (**Fig. 2**). This limited response could be linked to the early time point studied. In the current dataset, cisplatin was given at P21– 24 and OHCs were collected at P25, while cisplatin can be retained for a long time in the cochlea after treatment and can continuously damage the auditory system (Breglio et al., 2017). Our previous analysis also identified that, at this early stage, cisplatin induced transcriptional dysregulation across mitochondrial redox homeostasis, rather than cell death pathways such as apoptosis and ferroptosis. A previous in-vitro study in HEI-OC1 cells showed that cisplatin induces a time-dependent transcriptional change, with patterns of gene up/downregulation varying over time (Previati et al., 2004). Therefore, the observed changes in OHC functional transcripts in the current study reflect an early cellular response period, and later time points may show different expression patterns.

Overall, our findings show the context-dependent effects of honokiol on genes involved in OHC physiological function. The downregulation of MET-associated genes in the combination treatment group points to a potential mechanism that decreases cisplatin uptake, although its functional significance is yet to be investigated.

## Data Availability

The RNA-seq data in this work are available in the NCBI Gene Expression Omnibus under GSE328003. No new sequencing data were generated. Summaries for differential expression and interaction results are available in Supplementary Table 1.

## Acknowledgments

This work was supported by the Northwestern University NUSeq Core Facility.

## Author contributions

X.T. and C.K. conceived the research idea. X.T. acquired the funding. C.K. performed data analysis. C.K. and X.T. interpreted the results. C.K. and X.T. drafted the manuscript. All authors read and approved the final manuscript.

## Ethics declarations

No animal experiments were performed for the present study.

## Supplementary Information

Supplementary Table 1. Expression and interaction analysis of genes involved in outer hair cell function

